# Mixing patterns of HIV transmission among men who have sex with men in the United Kingdom

**DOI:** 10.1101/342774

**Authors:** Stéphane Le Vu, Oliver Ratmann, Valerie Delpech, Alison E Brown, O Noel Gill, Anna Tostevin, David Dunn, Christophe Fraser, Erik M Volz, on behalf of the UK HIV Drug Resistance Database^1^

## Abstract

**Background:** Near 60% of new HIV infections in the United Kingdom are estimated to occur in men who have sex with men (MSM). Patterns of mixing between different risk groups of MSM have been suggested to spread the HIV epidemics through age-disassortative partnerships and to contribute to ethnic disparities in infection rates. Understanding these mixing patterns in transmission can help to determine which groups are at a greater risk and guide prevention.

**Methods:** We analyzed combined epidemiologic data and viral sequences from MSM diagnosed with HIV as of mid-2015 at the national level. We applied a phylodynamic source attribution model to infer patterns of transmission between groups of patients by age, ethnicity and region.

**Results:** From pair probabilities of transmission between 19 847 MSM patients, we found that potential transmitters of HIV subtype B were on average 5 months older than recipients. We also found a moderate overall assortativity of transmission by ethnic group and a stronger assortativity by region.

**Conclusions:** Our findings suggest that there is only a modest net flow of transmissions from older to young MSM in subtype B epidemics and that young MSM, both for Black or White groups, are more likely to be infected by one another than expected in a sexual network with random mixing.

## 1. Introduction

Men who have sex with men (MSM) account for 40% of new diagnoses in Europe [1], and in the United Kingdom (UK), nearly sixty percent of new HIV infections are estimated to occur in MSM, although there is a recent sign of decline in transmission particularly recorded in London [2]. It has been estimated that the largest contribution to transmission in the UK is attributable to young HIV positive MSM (in this case less than 35) [3]. More generally, since the early work from Morris et al. [4], it has been suggested that young MSM having sex with older partners represent a significant driver of the epidemic in North America [5]. This disassortative age mixing pattern has also been studied in interaction with mixing in ethnicity [6, 7] or to illuminate disparity in HIV prevalence by ethnicity, which is a common feature of UK and US epidemics [8, 9].

Studies based on partnership data have shown that having an older partner increases the risk of infection [10, 11]. This result is also supported by analyses of transmission networks based on phylogenetic clustering [12, 13]. But clustering of genetically similar viruses is influenced by time since infection when patients are sampled, which is confounded by patients’ age as well as CD4 and clinical stage of infection. Phylogenetic clustering is also confounded by the fraction of infected persons sampled, which makes direct inference of transmission patterns difficult using genetic clustering [14–16].

When clustering is observed between older and younger MSM, it is suggestive of a flow from old to young because of higher prevalence in older MSM [12]. But, the direction of putative transmission events cannot be resolved by pairwise genetic distance alone, and it is not possible to estimate flows of transmission between age groups based on clustering observations. In this study, we applied a phylogenetic source attribution (SA) method that infers the probability of potential transmission (infector probability) between pairs of patients among approximately 20 000 MSM diagnosed in the UK with available genetic sequences [17]. Source attribution methods based on consensus pol-sequence data can not be used to infer transmission pairs with high confidence, but can provide useful insights when studied in aggregate over thousands of putative transmission pairs. In general, direction of transmission can not be inferred from consensus HIV sequence data, but in combination with clinical stage of infection at time of sequencing, directionality can be inferred probabilistically in some cases, as when for example a patient with chronic infection is linked to a patient with early infection. By combining phylogenetic analysis of consensus-pol sequences with stage of infection data and independent estimates of incidence and prevalence in the population, we are able to quantify potentially imbalanced transmission patterns between different risk groups including different age groups.

We used HIV-1 sequences routinely collected for drug resistance testing, patient-level data informative of the time since infection to account for biased sampling, and epidemiological estimates of background prevalence and incidence to account for potentially unsampled individuals that could be the sources of infection. In this study, by using estimates of transmission pair probabilities, our objective was to reveal patterns of transmission in men who have sex with men according to age, ethnicity, and geography. In particular, we searched for evidence of source-sink relationships in transmission patterns between age groups and examined the hypothesis that there is a net flow of transmissions from old to young MSM.

## 2. Materials and Methods

### 2.1. Data

We used partial HIV-1 pol sequences collected in the UK HIV Drug Resistance Database (http://www.hivrdb.org.uk/) linked with characteristics of patients newly diagnosed with HIV from the UK Collaborative HIV Cohort study database and the national HIV/AIDS Reporting System database [18], as of end of August 2016. The data were fully anonymised.

We analyzed adult patients reported as MSM; infected by HIV-1 subtype A1, B, C or CRF-02AG (the 4 most represented subtypes); and having a nucleotide sequence while treatment naive. The first sequence per patient with length > 950 nucleotides was included. CD4 count values closest to and within a maximum of 1 year of the date of sequence sampling were used to define 5 stages of infection, comprising early HIV infection (stage 1) and 4 stages of declining CD4 with thresholds at 500, 350 and 200 cells/mm3 [19]. In our sample, 81% of patients had a CD4 count information. A positive result from the avidity-based recent infection testing algorithm (RITA) led to classifying a patient as at stage 1. Results of RITA at diagnosis were available as of 2009, and from this year were informed for 46% of patients.

Age of patients was categorized in quartiles of age at the date of resistance testing. Ethnicity categories were grouped in 7 classes: White; Black Caribbean; Black African; Other or unspecified black; Indian, Pakistani or Bangladeshi (South Asian); Other Asian or Oriental, Other and mixed. Regions of diagnosis were categorized in 5 classes: London; South of England; Midlands and East of England; North of England; Northern Ireland, Scotland and Wales. In analyses of assortativity, unknown category was treated as missing data.

### 2.2. Sequence processing

Partial HIV-1 pol sequences were sampled from 1997 to July 2015 with a majority obtained after 2009. Sequences were aligned with MAFFT version 7 after adding 1780 international sequences from the Los Alamos HIV sequence database (http://www.hiv.lanl.gov/). The international sequences were added to infer importation of viral lineages, and were selected in a BLAST search that identified for each of the UK sequences the sequence with highest similarity (using bitscore) in the Los Alamos database. Drug resistance mutation sites as listed in the 2015 update from the IAS-USA were stripped from the alignment. Subtypes were determined with REGA version 3 and all further analyses were stratified by subtype.

### 2.3. Phylogenetic analysis

Phylogenetic trees were constructed with ExaML by maximum likelihood based inference with a gamma distribution model for rate heterogeneity among sites [20]. One hundred bootstrap replicates of each tree were computed to account for phylogenetic uncertainty. Trees were rooted at outgroup sequences that were added to each subtype alignment prior to tree reconstruction.

We calculated root-to-tip distance and regressed distance by time from MRCA to sample. By iterations of Grubb’s algorithm, we identified on overall 0.3% sequences as outliers in terms of divergence time and evolutionary rate. We applied least-square dating algorithm [21] on rooted trees and sampling times to estimate the substitution rate and dates of ancestral nodes.

We analyzed separately the 4 main subtypes to account for different evolutionary rates. Fitch algorithm was used to reconstruct ancestral host status and determine distinct clades of virus transmitted in the UK. The dated subtype B phylogeny comprised 18,484 taxa and for computational reasons was split into subtrees of <1000 sequences for further analyses.

### 2.4. Probabilistic source attribution

We applied a phylogenetic source attribution method that uses a population-genetic model to derive probabilities that a given individual (donor) is the source of infection for another individual (recipient) in the sample. These probabilities, termed *infector probabilities*, account for the epidemiological and sampling processes by incorporating into their calculation the time-scaled phylogeny, patient data on stage of infection, and population-level data on occurrence of infection [17]. The method was evaluated in a previous simulation study [16].

For population-level epidemic statistics, we used updated incidence estimates of CD4-based back-calculation method for MSM population and prevalence estimates of Bayesian multiparameter synthesis of surveillance data, as reported by Public Health England in 2017 [2]. To account for uncertainty in those input parameters, we randomly drew 5 pair values of incidence and prevalence per bootstrap replicates (2000 in total) from normal distributions inferred from the credible intervals of those estimates. Incidence and prevalence were assumed to be proportional across subtypes.

The source attribution method uses a continuous time Markov chain model to reconstruct the likely state of a lineage at the time of transmission given the CD4-stage of infection at time of sampling. The definition of stages of infection and progression rates were based on Cori et al. [19], as described in our previous analysis [16]. The method assumes that each infected patient corresponds to a single lineage of virus, ignoring multiple infections, and that internal nodes in the phylogeny correspond to a transmission event between hosts. To limit calculations to non-negligible pairing, only coalescent events within a limit of 20 years prior to sequence sampling were incorporated to compute infector probabilities.

### 2.5. Statistical procedures

Infector probabilities *W*_*ij*_ for each donor-recipient pair were averaged over all bootstrap replicates. To compare the mean age of donors and recipients we used a two-tailed paired weighted t-test, with pair-level infector probabilities as weights.

To characterize transmission patterns by patients’ covariates, we first computed a symmetric mixing matrix *M* as the normalized sum of infector probabilities representing aggregated number of transmissions between category *k* (*k* = 1,…,*m*) of recipients and category *l* (*l* = 1,…,*m*) of donors defined by age, ethnicity and region of diagnosis 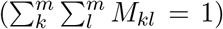. We then calculated 3 types of output matrices: (1) 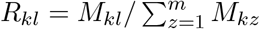, representing the conditional probability for a recipient in category *k* of being infected by a donor in category *l*; (2) 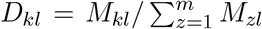, representing the conditional probability for a donor in category *l* of having transmitted to a recipient in category *k*; and (3) *A* = (*M* − *E*)/*E*, the assortativity matrix representing excessive transmission between categories of donors and recipients relative to random allocation. The matrix *E* has elements *E*_*kl*_ = Σ_*k*_ *M*_*kl*_ ⊗ Σ_*l*_ *M*_*kl*_/Σ_*k*_Σ_*l*_ *M*_*kl*_, and represents the expected values in the absence of preferential mixing [22]. Matrix *E* allows the calculation of Newman’s assortativity coefficient *r* = (Tr(*M*) − Tr(*E*))/(1 − Tr(*E*)). The coefficient ranges from −1 to 1, where *r* = 0 when there is no assortative mixing, *r* = 1 when there is perfect assortativity (every link connects individuals of the same type), and some negative value −1 ≤ *r* < 0 for a perfectly disassortative network (the lower bound depending on number of categories and density of subgraphs in each category). In all matrix-type figures, we represent transmission going from donors in columns to recipient in rows.

## 3. Results

### 3.1. Characteristics of the study population

The demographic and geographic composition of the 19,847 HIV-1 partial pol sequences from treatment naive patients diagnosed in the UK is described in table 1. Most gay and bisexual men diagnosed in the UK were infected with subtype B (93%). Therefore the patterns of transmission inferred from reconstructed phylogeny of subtype B sequences are largely dominating that of all MSM patients. Patients infected with non-B subtype were on average sampled later (median year of 2008 for subtype B, 2009 for subtypes A1 and C, and 2011 for CRF02AG) and were on average younger (median age of 35 for subtype B, 34 for subtypes A and C, and 32 for CRF02AG).

**Table 1:**
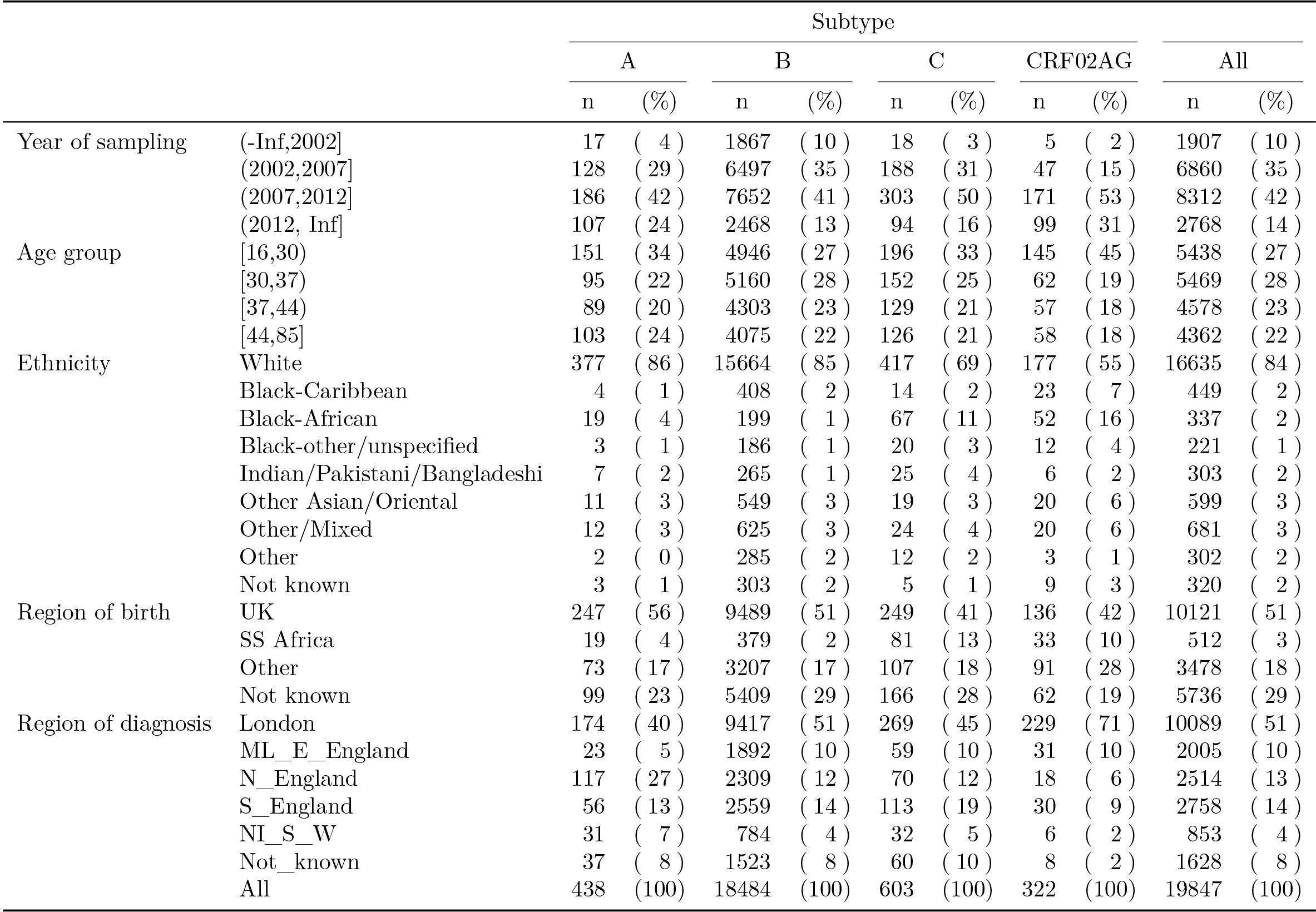
Characteristics of the study population

In terms of ethnicity, the majority (84%) of patients were white persons. Patients infected with C or CRF02AG were more commonly of non-white ethnicity: Black-African for 11% and 16% and from other non-white ethnicity for 19% and 26% respectively.

In terms of geography, half of subtype B and 71% of subtype CRF02AG sequences were sampled in Greater London. Apart from London, subtype A was especially prevalent in North of England (27%).

### 3.2. Infector probabilities

Across 100 bootstrap tree replicates for each subtype, we computed infector probabilities for on average 554 514 potential transmission pairs of 14 603 patients (cf. table 2). Given the *n* by *n* matrix of probabilities that a patient *i* transmitted to a patient *j*, the sum Σ_*i*_ *W*_*ij*_ represents the probability that the infector of *j* is in the sample. This quantity, denoted ‘in-degree’ in table 2, indicates that on average 36.6% (95%CI[35.2 - 38.0]) of potential donors are included in our sampled population. Our estimates of in-degrees were moderately influenced by the variation in inputs of background incidence and prevalence, with lower incidence (or higher prevalence) increasing average in-degrees as the probability of an unsampled intermediary transmitter is decreased (cf. figure S1).

**Table 2:** Phylogenetic reconstruction and source attribution results. *Global sequences* are unique sequences from Los Alamos HIV sequence database matching UK sequences from a BLAST search. *Outliers* are UK sequences identified as outliers in root-to-tip regression.

| Subtype | A | B | C | CRF02AG | All |
| --- | --- | --- | --- | --- | --- |
| Global sequences (n) | 199 | 831 | 612 | 138 | 1780 |
| Outliers (n) | 6 | 163 | 7 | 74 | 250 |
| Median TMRCA (year) | 1951 | 1966 | 1961 | 1975 |  |
| Donors/recipients (mean) | 337 | 13665 | 346 | 255 | 14603 |
| Potential pairs (mean) | 19818 | 521811 | 6350 | 6535 | 554514 |
| Mean in-degree (%) | 39.4 | 36.7 | 32.6 | 28.9 | 36.6 |

### 3.3. Age difference between donors and recipients

Table 3 shows the mean difference in age between donors and recipients, weighted by infector probabilities. A significant difference is only detectable for subtype B, donors being on average 5 months older than recipients. For subtype B, most transmission pairs in our sample involved individuals less than 30 years old (figure 1M). The largest proportion (46%) of infection acquired by young individuals was attributable to individuals in the same age category (figure 1R). And a strong assortativity in transmission mixing is seen in this youngest age category, indicating that young MSM are preferentially infected by young MSM. This preferential mixing is also seen in among individuals over 44 years. The overall assortativity coefficient was moderate with *r* = 0.16. Similar transmission patterns between age groups were observed for subtypes A and C (table S2). However transmission of subtype CRF02AG was characterized by a strong assortativity mostly in the oldest age category but more intergenerational mixing between other categories(figure S2A). Despite the lack of significant difference in average age of donors relative to recipient shown previously for subtype CRF02AG, the most probable infector for individuals from intermediate age quartiles (30-36 and 37-43) was younger (less than 30) (figure S2R).

**Table 3:**
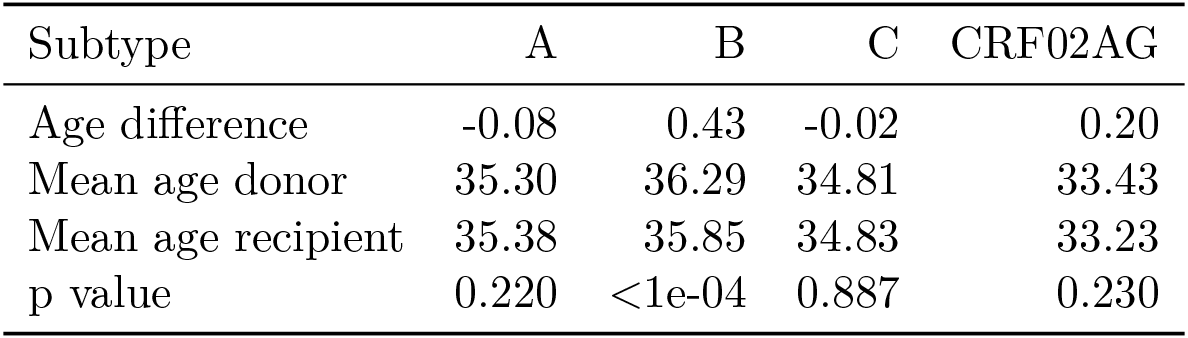
Difference between age of donor and age of recipient

| Subtype | A | B | C | CRF02AG |
| --- | --- | --- | --- | --- |
| Age difference | -0.08 | 0.43 | -0.02 | 0.20 |
| Mean age donor | 35.30 | 36.29 | 34.81 | 33.43 |
| Mean age recipient | 35.38 | 35.85 | 34.83 | 33.23 |
| p value | 0.220 | <1e-04 | 0.887 | 0.230 |

**Figure 1:**
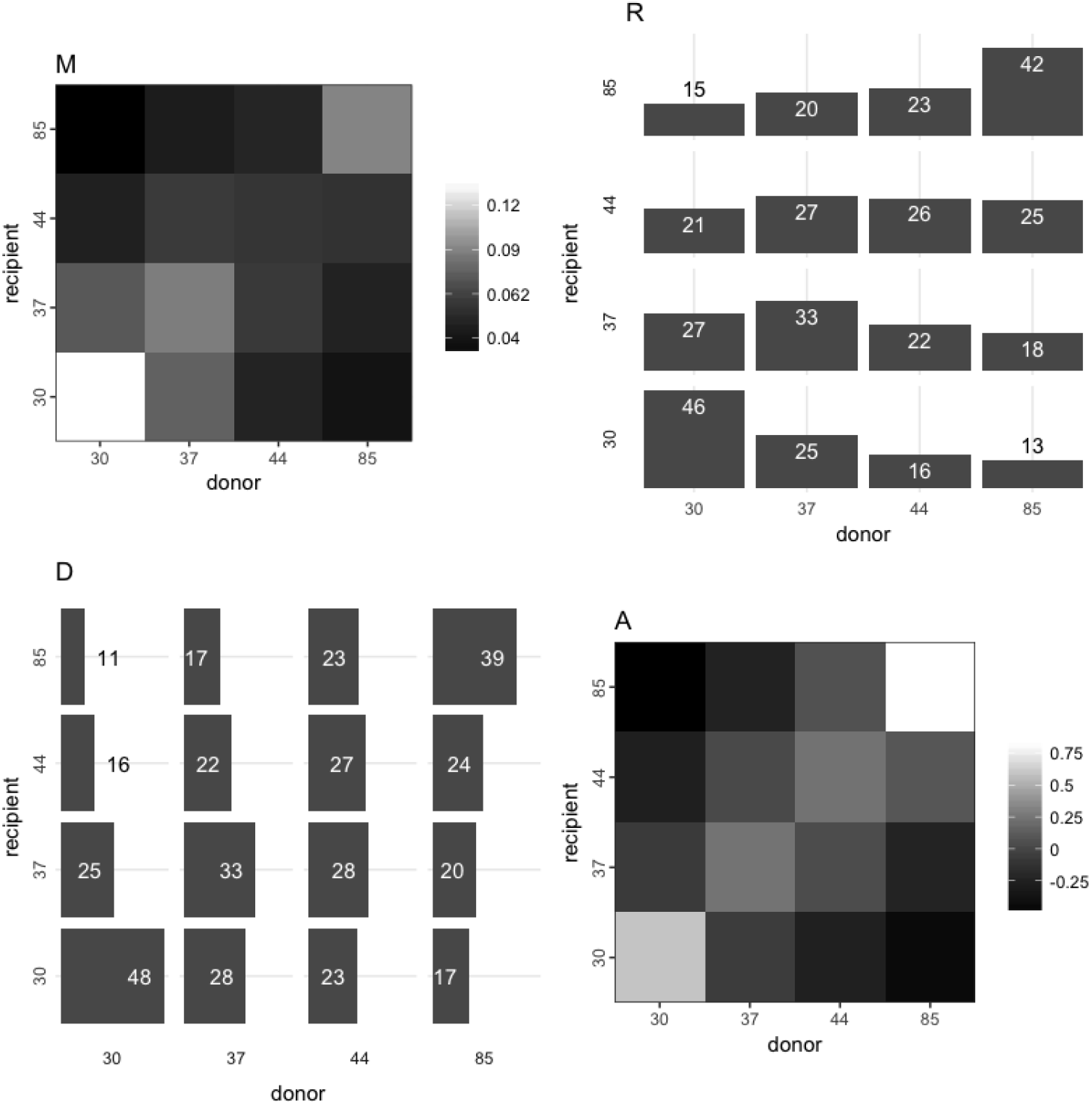
Patterns of transmission of HIV subtype B by age in quartiles. The four graphics depict transmission from donor categories in column to recipient categories in row (from x-axis to y-axis). Axes labels represent the upper bound of quartiles of age. **M**: Each cell represents the proportion of overall transmissions from one category to an other, with higher proportion in lighter shade, *i.e.* the highest amount (14%) of transmissions involved donors and recipients both aged less than 30. **R**: Each row represents the probability distribution for a given age category of recipients of having been infected by donors by age, *i.e.* 25% of recipients less than 30 years old were infected by donor aged 30-36. **D**: Each column represents the probability distribution for a given age category of donors of having transmitted to recipients by age, *i.e.* 28% of donors aged 37-43 infected recipients aged 3036. **A**: The assortativity matrix indicates that, relative to random mixing more transmissions occurred within the same age category, particularly for the oldest and youngest. Assortativity coefficient *r* = 0.16.

### 3.4. Transmission by ethnicity

The vast majority (85%) of MSM infected with subtype B viruses were of white ethnicity. We estimated that 82% of all transmissions in our sample occurred between white individuals, and that recipients of all ethnicities had a majority of white donors. The probability of having been infected by a white individual was 92% for whites, 77% for Indian/Pakistani or Bengladeshi, 75% for other Asians, 55% for Black Africans and 54% for Blacks Caribbeans. Conversely, a majority of transmission originating from donors of any ethnic group was estimated to affect white recipients. Figure 2a shows the level of assortativity in transmission of subtype B viruses between ethnic groups. Inter-ethnic transmission (cumulated pair probabilities outside the diagonal) represented 17% on overall and 58% when excluding the white category. Overall assortativity was moderate (*r* = 0.17) but a preferential mixing was especially observed within and between all black ethnic groups and within the South Asian group.

**Figure 2:**
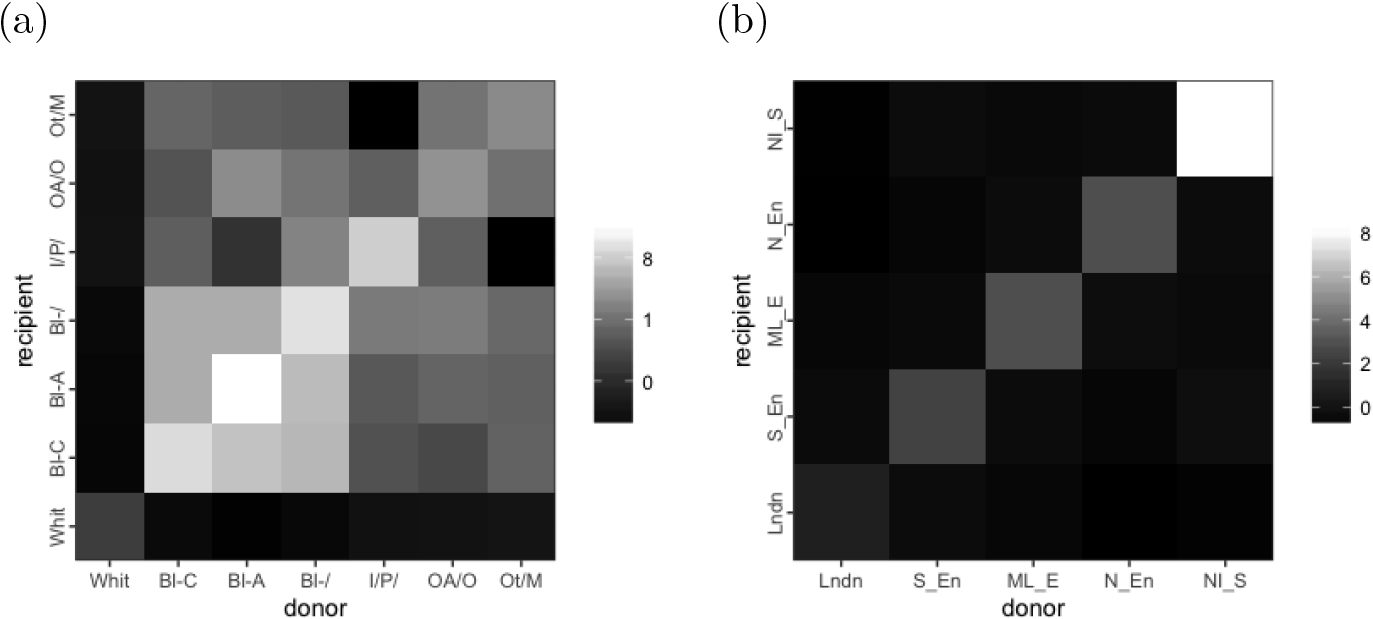
Assortativity in transmission of HIV-1 subtype B by ethnicity and region of diagnosis. Lighter shades represent higher assortativity. **(a)**: Ethnicities: White; Black Caribbean (Bl-C); Black African (Bl-A); Other or unspecified black (Bl-); Indian, Pakistani or Bangladeshi (I/P/); Other Asian or Oriental (OA/O), Other and mixed (Ot/M). Assortativity coefficient *r* = 0.17. **(b)**: Regions: London, South of England; Midlands and East of England; North of England; Northern Ireland, Scotland and Wales. Assortativity coefficient *r* = 0.56.

We estimated the probability of transmission of subtype B viruses between young (<30) and older MSM (30+) either from white or black ethnicity (figure 3). The relative excess of transmission within age categories observed previously is observed for both white and black ethnicities, and overall assortativity by age was similar (*r* = 0.25 for white and 0.28 for black). However, for a given older MSM the probability of transmitting to a young MSM was higher in black (39%) than in white ethnic group (22%).

**Figure 3:**
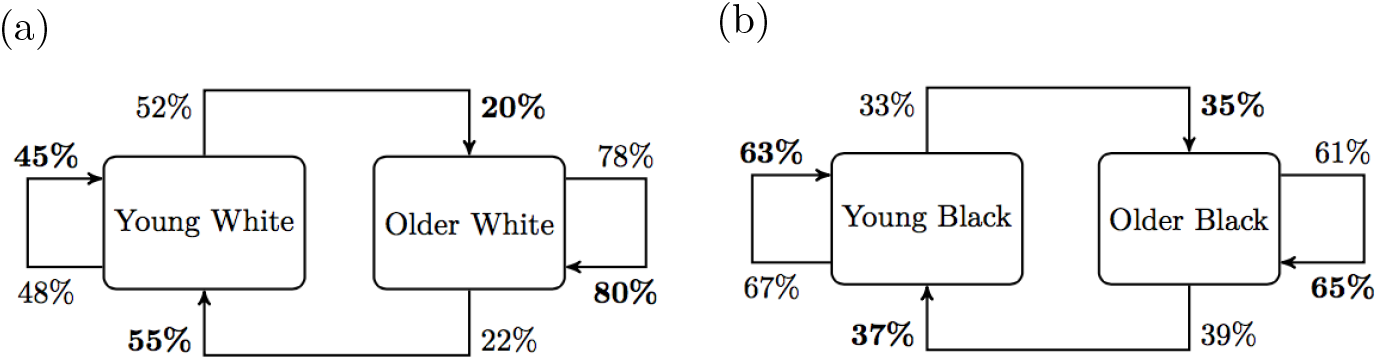
Patterns of transmission of HIV-1 subtype B between young MSM (less than 30) and older MSM by ethnicity: (a) White, (b) Black (including Black Africans, Black Carribean and other and unspecified Black). Percentages represent conditional probability of transmitting to recipient type per donor type (normal font) and of acquiring infection from donor type per recipient type (bold font).

### 3.5. Transmission by geographical region

Analyses of transmission by region show the largest level of assortativity, indicating a overall strong spatial structure of the epidemics (see figure 2b). Assortativity coefficients were 0.56 for subtype B and 0.49 for subtype CRF02AG. For those two subtypes, figure 4 shows the probability for a donor in a given region to transmit to a recipient of each respective region. For subtype B (left), the majority of transmissions (at least 60%) occur within the same region but donors from every region contributed to infections diagnosed in London (10% for North of England, Northern Ireland, Scotland and Wales, 20% for the Midlands and East England, and 30% for the South of England). For subtype CRF02AG, there was a higher probability for donors from North of England (60%) or Northern Ireland, Scotland and Wales (70%) to infect recipients in London than individuals within the same region.

**Figure 4:**
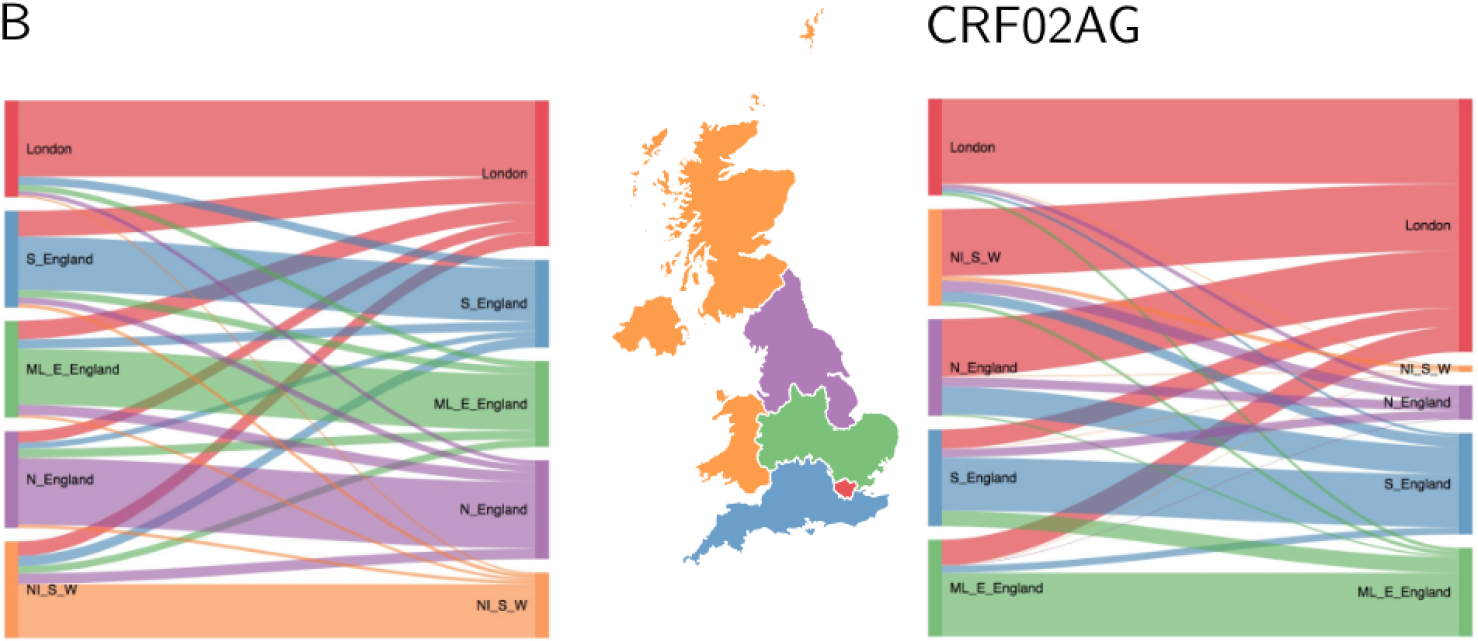
Patterns of transmission of HIV-1 subtype B (left) and CRF02AG (right), by geography. Each flow diagram, obtained from *D* matrix described in Methods section, has connections proportional to the probability of transmission from a donor given his region (left side) to recipients from respective regions (right side). The map is colored by groups of region of diagnosis: London, South of England (S_England); Midlands and East of England (ML_E_England); North of England (N_England); Northern Ireland, Scotland and Wales (NI_S_W).

## 4. Discussion

The objective of this study was to describe patterns of HIV transmission between age, ethnicity and geographical categories in the United Kingdom. We used a phylodynamic inference based on sequences collected among diagnosed MSM which accounts for incomplete sampling and stage of infection at sampling time. By modelling an epidemic process that is compatible with the evolution of transmitted viruses and epidemiological surveillance data, we characterized past transmission events among nearly 15 000 MSM patients at the national level.

Pair probabilities averaged over phylogenies and aggregated by age groups indicated a modest overall net flow of transmission from older to young MSM. This result is compatible with other studies reporting co-clustering of young and older patients [12, 13] as we do not observe pure assortative mixing, with probable transmission occurring in both directions across age groups. But our results indicate that on average, flow from old to young is mostly compensated by the transmission from young to old (cf. figure 1). And when the flow is imbalanced, as for transmission of subtype B viruses, the difference is small.

We observed an overall preferential mixing in transmission by age with greater assortativity both in the youngest and oldest age groups and more random mixing in intermediate age groups. Understanding age mixing patterns in transmission can help to determine which groups are at a greater risk and potentially guide public health interventions [23]. Our findings confirm that young MSM infect one another more than expected by random mixing which supports the idea that prevention benefit could be enhanced by focusing on this small group [24]. This result also corroborates the observation of recent clusters of young MSM sustaining the epidemic in the Netherlands [25].

We showed an overall preferential pairing by ethnicity in conjunction with an important mixing between white men and men from each other ethnicity. It can be explained by the overwhelming proportion of white men in the population. But in non-white groups, more than a half of transmission was inter-ethnic, revealing that a substantial amount of transmission has occurred between ethnic groups among MSM. A similar pattern for sexual partnership between ethnic groups was reported in Britain [8]. Although we found a relatively higher assortativity among black MSM in general and a non negligible mixing between black ethnic groups from different origins (African, Caribbean and other), HIV transmission seems less assortative among black MSM in the UK than it is in the USA [26].

We assessed whether intergenerational transmission was different in white and black MSM and found a similar level of age assortativity in both groups. Therefore as others in the US context [7] we did not find support in our findings to explain a disparity in HIV prevalence by age mixing [5, 6].

Finally, we found a strong geographical structure for the epidemics among MSM, with region of diagnosis as the variable associated with the highest level of assortativity. However, region of diagnosis can be different than the region of residency or of actual transmission which is not known, which may lead to an underestimation of the true level of geographical structure.

Several potential limitations of our study relates to the assumptions of the phylogenetic inference and source attribution method. First, as stated in methods section, the source attribution method neglects some effects of within-host evolution which can cause discordance between phylogenies and transmission trees [27]. This approximation is reasonable if within-host evolution generates coalescence time considerably shorter than between hosts at the population level. Secondly, we incorporated crude estimates of incidence and prevalence in the inference of infector probabilities. These were assumed constant over the period and proportional across subtypes. However variation of these inputs within credible limits had limited impact on average infector probabilities (figure S1). Third, the direction in transmission was derived from CD4 count and RITA result data that were partially complete.

Nevertheless, our analysis aimed to improve the use of phylogenetic information relative to genetic clustering in two ways. First, by providing a rough measure of transmission probability which unlike linkage into clusters can indicate a directionality and gives more weight to pairs with higher credibility. Notably, output matrices and patterns between groups would be symmetrical if based on clustering. Secondly, by correcting for biases stemming from incomplete sampling of the infected host population. Lastly, the source attribution method was fast to compute and scaled easily to phylogenies based on many thousands of sequences.

Future directions for this work include applying the analysis to the heterosexual population, where phylogenetic information could contribute to assess age disparity in mixing across gender. [28, 29]. Another direction would be to use methods exploiting next-generation sequencing, that account for within-host evolution and enhance resolution in identifying transmission [27, 30].

In conclusion, this study has leveraged available patients data and viral sequences to provide evidence of assortativity in HIV transmission by age, ethnicity and geography. Understanding these patterns of transmission is important to modelling the impact of intervention strategies.

## Acknowledgements

This work was supported by the National Institute for Health Research (NIHR) Health Protection Research Units in Modeling Methodology and Sexually Transmitted Infections (HPRU-2012-10080). E.M.V. is supported by the National Institutes of Health (R01AI087520). O.R. and C.F. are supported by Bill & Melinda Gates Foundation: Phylogenetics Networks to Address Transmission of HIV (OPP1084362). A.T. is supported by UK HIV Drug Resistance Database grant from the Medical Research Council (164587). We thank the Imperial College High Performance Computing Service (doi: 10.14469/hpc/2232).

## Supplementary material for the article “Mixing patterns of HIV transmission among men who have sex with men in the United Kingdom”

**Figure S1:**
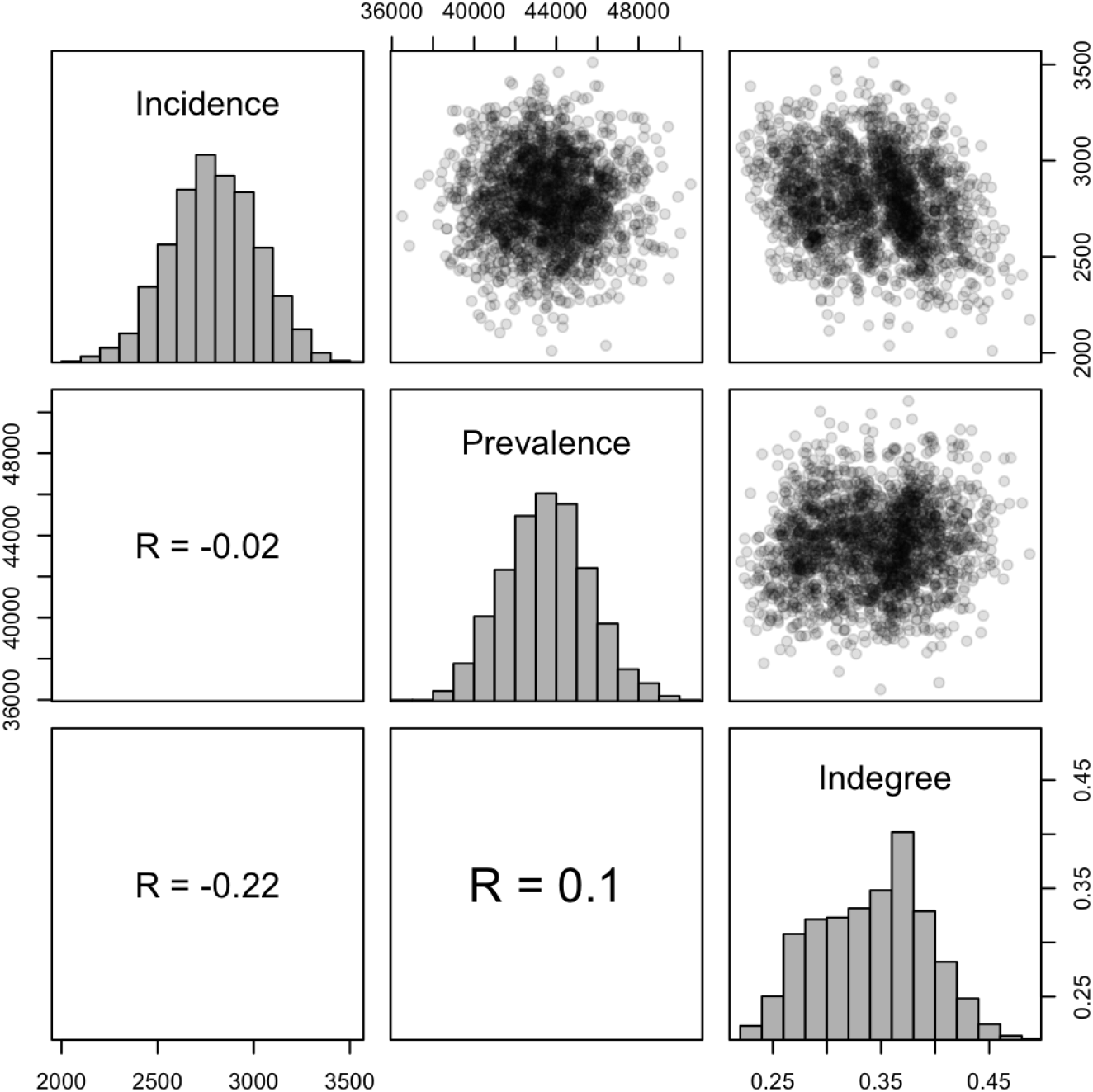
Influence of variation in incidence and prevalence inputs on mean indegree. Incidence (number of new infections per year) and prevalence (number of persons living with HIV) were drawn independently 5 times per each of 100 phylogenetic trees for subtype B transmission. Under the source attribution model, the in-degree is the sum of infector probabilities incoming to an individual. It also represents the probability that its infector is in the sample. The mean in-degree tends to increase as prevalence is increased or as incidence is lowered, as it decreases the probability of an unsampled source case. Both variations have a limited impact on the average in-degree estimates as indicated by correlation coefficient R.

**Figure S2:**
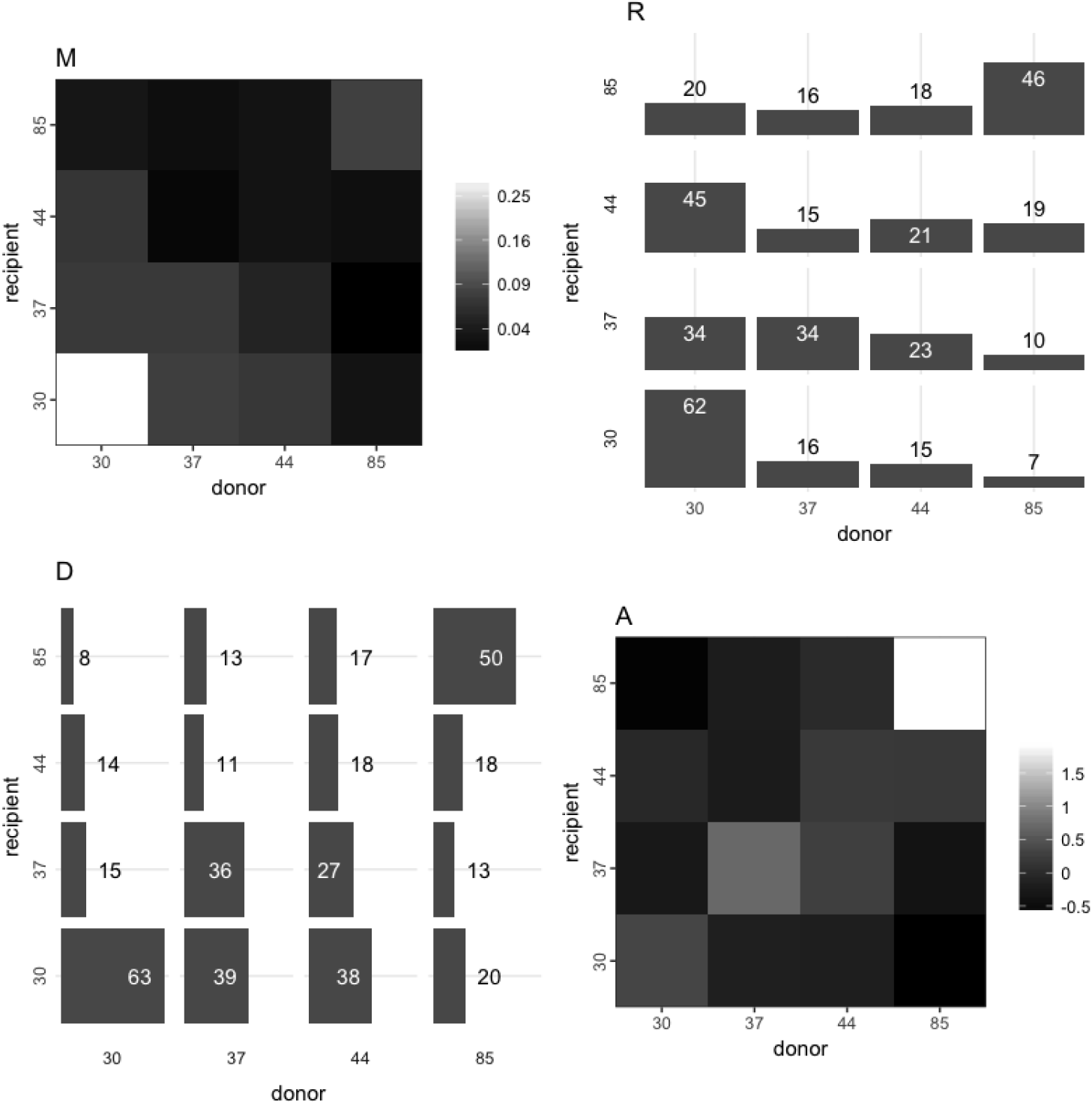
Patterns of transmission of HIV-1 subtype CRF02AG by age. See reading notes from figure 1 in main text.

**Figure S3:**
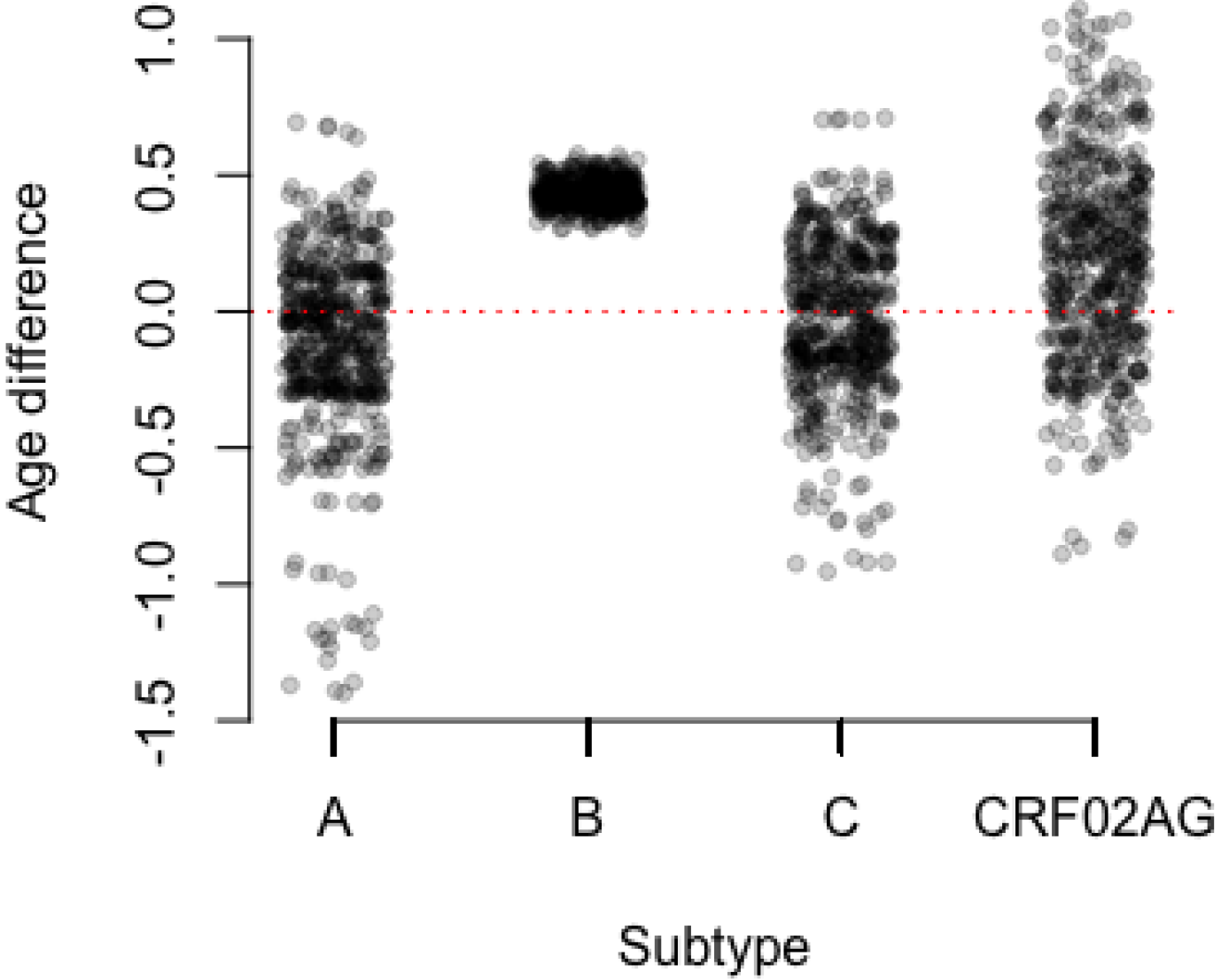
Age difference between donors and recipients by subtype across 100 bootstrap replicates. Differences are in year. Dotted red line indicates no difference.

**Table S1:**
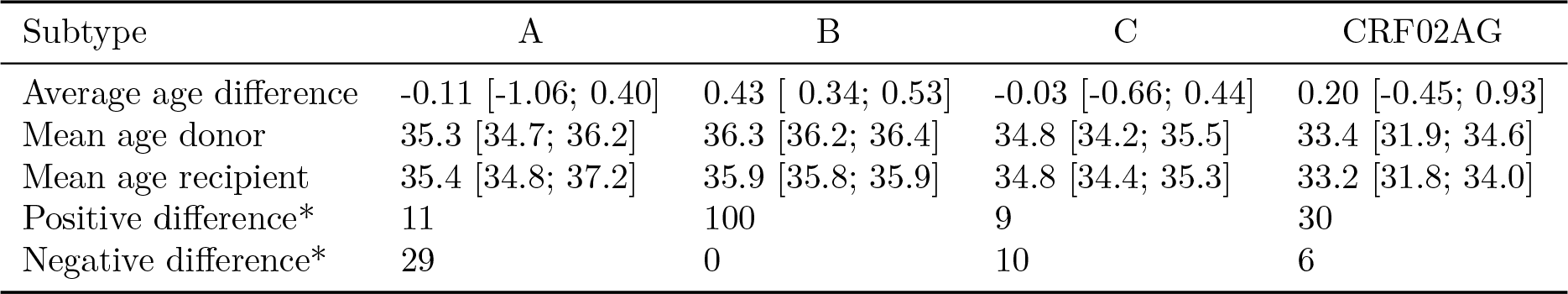
Difference in year between age of donor and age of recipient, across 100 bootstrap replicates. *Number of p-values < 0.05 for two-tailed weighted t-test of the age difference.

| Subtype | A | B | C | CRF02AG |
| --- | --- | --- | --- | --- |
| Average age difference | -0.11 [-1.06; 0.40] | 0.43 [0.34; 0.53] | -0.03 [-0.66; 0.44] | 0.20 [-0.45; 0.93] |
| Mean age donor | 35.3 [34.7; 36.2] | 36.3 [36.2; 36.4] | 34.8 [34.2; 35.5] | 33.4 [31.9; 34.6] |
| Mean age recipient | 35.4 [34.8; 37.2] | 35.9 [35.8; 35.9] | 34.8 [34.4; 35.3] | 33.2 [31.8; 34.0] |
| Positive difference* | 11 | 100 | 9 | 30 |
| Negative difference* | 29 | 0 | 10 | 6 |

**Table S2:** 95% confidence intervals for proportion of transmission by age quartiles. M, R and D columns refer to the matrices described in section 2.5 and represented in figure 1. Column labels for donors and row labels for recipients represent the upper bound of quartiles of age.

| Subtype | M |  |  |  | R |  |  |  | D |  |  |  |
| --- | --- | --- | --- | --- | --- | --- | --- | --- | --- | --- | --- | --- |
|  | 30 | 37 | 44 | 85 | 30 | 37 | 44 | 85 | 30 | 37 | 44 | 85 |
| **A** |  |  |  |  |  |  |  |  |  |  |  |  |
| 85 | 03;08 | 03;05 | 06;09 | 06;09 | 14;27 | 12;20 | 24;34 | 26;40 | 09;25 | 15;23 | 27;46 | 31;43 |
| 44 | 03;04 | 02;04 | 04;07 | 06;09 | 16;23 | 11;20 | 21;34 | 33;45 | 08;12 | 10;18 | 20;29 | 28;39 |
| 37 | 07;09 | 04;08 | 02;05 | 03;04 | 33;45 | 19;35 | 08;25 | 12;20 | 19;25 | 19;35 | 10;23 | 11;19 |
| 30 | 15;23 | 07;11 | 04;06 | 03;05 | 47;60 | 19;28 | 10;16 | 07;15 | 44;61 | 35;48 | 19;28 | 13;21 |
| **B** |  |  |  |  |  |  |  |  |  |  |  |  |
| 85 | 03;03 | 04;05 | 05;05 | 09;09 | 14;16 | 20;21 | 22;23 | 41;43 | 11;12 | 16;17 | 22;24 | 38;40 |
| 44 | 04;05 | 06;06 | 05;06 | 05;06 | 20;22 | 26;28 | 25;27 | 24;26 | 15;16 | 21;23 | 26;28 | 23;24 |
| 37 | 07;07 | 09;09 | 06;06 | 05;05 | 26;28 | 32;34 | 21;23 | 17;19 | 24;25 | 32;34 | 27;29 | 20;21 |
| 30 | 14;14 | 07;08 | 05;05 | 04;04 | 45;47 | 24;26 | 15;16 | 12;14 | 47;49 | 28;29 | 22;24 | 16;18 |
| **C** |  |  |  |  |  |  |  |  |  |  |  |  |
| 85 | 01;03 | 03;05 | 03;04 | 05;09 | 07;16 | 17;30 | 17;28 | 34;51 | 03;07 | 10;18 | 14;22 | 30;49 |
| 44 | 05;07 | 05;07 | 07;10 | 03;06 | 20;27 | 21;30 | 28;38 | 13;22 | 13;19 | 19;27 | 36;47 | 19;31 |
| 37 | 08;11 | 05;09 | 04;06 | 03;05 | 31;43 | 21;33 | 16;24 | 11;20 | 23;31 | 21;31 | 21;32 | 16;30 |
| 30 | 17;21 | 08;13 | 02;04 | 01;03 | 50;61 | 25;37 | 06;11 | 04;09 | 48;57 | 32;44 | 10;19 | 08;17 |
| **CRF02AG** |  |  |  |  |  |  |  |  |  |  |  |  |
| 85 | 02;05 | 02;04 | 01;04 | 05;11 | 13;30 | 10;24 | 11;26 | 31;58 | 05;10 | 09;19 | 08;23 | 35;61 |
| 44 | 05;08 | 01;04 | 02;04 | 01;04 | 33;53 | 09;23 | 16;30 | 10;25 | 09;18 | 07;19 | 13;26 | 10;27 |
| 37 | 05;09 | 05;09 | 04;07 | 01;03 | 27;41 | 28;41 | 17;29 | 06;15 | 12;19 | 28;47 | 20;37 | 08;19 |
| 30 | 25;34 | 05;10 | 05;09 | 02;04 | 55;69 | 11;23 | 10;19 | 04;09 | 56;68 | 29;49 | 29;46 | 13;28 |

## References

[1] European Centre for Disease Prevention and Control, WHO Regional Office for Europe, HIV/AIDS surveillance in Europe 2017 -2016 data, Tech. rep., ECDC, Stockholm (2017). URL http://ecdc.europa.eu/en/publications-data/hivaids-surveillance-europe-2017-2016-data

[2] A. E. Brown, P. Kirwan, C. Chau, J. Khawam, O. N. Gill, V. C. Delpech, Towards elimination of HIV transmission AIDS and HIV related deaths in the UK -2017 report, Tech. rep., Public Health England (2017).URL https://www.gov.uk/government/uploads/system/uploads/attachment_data/file/662306/Towards_elimination_of_HIV_transmission_AIDS_and_HIV_related_deaths_in_the_UK.pdf

[3] N. Punyacharoensin, W. J. Edmunds, D. De Angelis, V. Delpech, G. Hart, J. Elford, A. Brown, N. Gill, R. G. White, Modelling the HIV epidemic among MSM in the United Kingdom: Quantifying the contributions to HIV transmission to better inform prevention initiatives, AIDS (London, England) 29 (3) (2015) 339–349. doi:10.1097/QAD.0000000000000525.

[4] M. Morris, J. Zavisca, L. Dean, Social and sexual networks: Their role in the spread of HIV/AIDS among young gay men, AIDS education and prevention: official publication of the International Society for AIDS Education 7 (5 Suppl) (1995) 24–35.URL https://www.ncbi.nlm.nih.gov/pubmed/8664095.

[5] B. J. Coburn, S. Blower, A Major HIV Risk Factor for Young Men Who Have Sex With Men Is Sex With Older Partners:, JAIDS Journal of Acquired Immune Deficiency Syndromes (2010) 1 doi:10.1097/QAI.0b013e3181d43999..

[6] M. Berry, H. F. Raymond, W. Mcfarland, Same race and older partner selection may explain higher Hiv prevalence among black men who have sex with men, Aids 21 (17) (2007) 2349–2350. doi:10.1097/QAD.0b013e3282f12f41.

[7] J. A. Grey, R. B. Rothenberg, P. S. Sullivan, E. S. Rosenberg, Disas-sortative Age-Mixing Does Not Explain Differences in HIV Prevalence between Young White and Black MSM: Findings from Four Studies, PLoS ONE 10 (6). doi:10.1371/journal.pone.0129877.

[8] R. Doerner, E. McKeown, S. Nelson, J. Anderson, N. Low, J. Elford, Sexual Mixing and HIV Risk Among Ethnic Minority MSM in Britain, AIDS and Behavior 16 (7) (2012) 2033–2041. doi:10.1007/s10461-012-0265-3.

[9] G. A. Millett, J. L. Peterson, S. A. Flores, T. A. Hart, W. L. Jeffries, P. A. Wilson, S. B. Rourke, C. M. Heilig, J. Elford, K. A. Fenton, R. S. Remis, Comparisons of disparities and risks of HIV infection in black and other men who have sex with men in Canada, UK, and USA: A meta-analysis, The Lancet 380 (9839) (2012) 341–348. doi:10.1016/ S0140-6736(12)60899-X.

[10] C. B. Hurt, D. D. Matthews, M. S. Calabria, K. A. Green, A. A. Adimora, C. E. Golin, L. B. Hightow-Weidman, Sex with Older Partners Is Associated With Primary HIV Infection Among Men Who Have Sex With Men in North Carolina:, JAIDS Journal of Acquired Immune Deficiency Syndromes (2010) 1 doi:10.1097/QAI.0b013e3181c99114.

[11] F. Jin, A. E. Grulich, L. Mao, I. Zablotska, M. O’Dwyer, M. Poynten, G. P. Prestage, Sexual partner’s age as a risk factor for HIV seroconversion in a cohort of HIV-negative homosexual men in Sydney, AIDS and behavior 17 (7) (2013) 2426–2429. doi:10.1007/s10461-012-0350-7.

[12] Y. O. Whiteside, R. Song, J. O. Wertheim, A. M. Oster, Molecular analysis allows inference into HIV transmission among young men who have sex with men in the United States, AIDS (London, England) 29 (18) (2015) 2517–2522. doi:10.1097/QAD.0000000000000852.

[13] E. Wolf, J. T. Herbeck, S. Van Rompaey, M. Kitahata, K. Thomas, G. Pepper, L. Frenkel, Phylogenetic Evidence of HIV-1 Transmission Between Adult and Adolescent Men Who Have Sex with Men, AIDS Research and Human Retroviruses 33 (4) (2016) 318–322. doi:10.1089/aid.2016.0061.

[14] E. M. Volz, J. S. Koopman, M. J. Ward, A. L. Brown, S. D. W. Frost, Simple epidemiological dynamics explain phylogenetic clustering of HIV from patients with recent infection, PLoS computational biology 8 (6) (2012) e1002552. doi:10.1371/journal.pcbi.1002552.

[15] A. F. Y. Poon, Impacts and shortcomings of genetic clustering methods for infectious disease outbreaks, Virus Evolution 2 (2) (2016) vew031. doi:10.1093/ve/vew031.

[16] S. Le Vu, O. Ratmann, V. Delpech, A. E. Brown, O. N. Gill, A. Tostevin, C. Fraser, E. M. Volz, Comparison of cluster-based and source-attribution methods for estimating transmission risk using large HIV sequence databases, Epidemics doi:10.1016/j.epidem.2017.10.001.

[17] E. M. Volz, S. D. W. Frost, Inferring the Source of Transmission with Phylogenetic Data, PLoS Computational Biology 9 (12). doi:10.1371/journal.pcbi.1003397.

[18] Public Health England, HIV and AIDS Reporting System (2018). URL https://www.gov.uk/government/collections/hiv-surveillance-data-and-management.

[19] A. Cori, M. Pickles, A. van Sighem, L. Gras, D. Bezemer, P. Reiss, C. Fraser, CD4+ cell dynamics in untreated HIV-1 infection: Overall rates, and effects of age, viral load, sex and calendar time, AIDS (London, England) 29 (18) (2015) 2435–2446. doi:10.1097/QAD.0000000000000854.

[20] A. M. Kozlov, A. J. Aberer, A. Stamatakis, ExaML version 3: A tool for phylogenomic analyses on supercomputers, Bioinformatics 31 (15) (2015) 2577–2579. doi:10.1093/bioinformatics/btv184.

[21] T.-H. To, M. Jung, S. Lycett, O. Gascuel, Fast Dating Using Least-Squares Criteria and Algorithms, Systematic Biology 65 (1) (2016) 82–97. doi:10.1093/sysbio/syv068.

[22] M. E. J. Newman, Mixing patterns in networks, Physical Review E 67 (2) (2003) 026126. doi:10.1103/PhysRevE.67.026126.

[23] A. Anema, B. D. L. Marshall, B. Stevenson, J. Gurm, G. Montaner, W. Small, E. A. Roth, V. D. Lima, J. S. G. Montaner, D. Moore, R. S. Hogg, Intergenerational sex as a risk factor for HIV among young men who have sex with men: A scoping review, Current HIV/AIDS reports 10 (4) (2013) 398–407. doi:10.1007/s11904-013-0187-3.

[24] E. M. Volz, S. Le Vu, O. Ratmann, A. Tostevin, D. Dunn, C. Orkin, S. O’Shea, V. Delpech, A. Brown, N. Gill, C. Fraser, UK HIV Drug Resistance Database, Molecular Epidemiology of HIV-1 Subtype B Reveals Heterogeneous Transmission Risk: Implications for Intervention and Control, The Journal of Infectious Diseases 217 (10) (2018) 1522–1529. doi:10.1093/infdis/jiy044.

[25] D. Bezemer, A. Cori, O. Ratmann, A. van Sighem, H. S. Hermanides, B. E. Dutilh, L. Gras, N. Rodrigues Faria, R. van den Hengel, A. J. Duits, P. Reiss, F. de Wolf, C. Fraser, ATHENA observational cohort, Dispersion of the HIV-1 Epidemic in Men Who Have Sex with Men in the Netherlands: A Combined Mathematical Model and Phylogenetic Analysis, PLoS Med 12 (11) (2015) e1001898. doi:10.1371/journal.pmed.1001898.

[26] A. M. Oster, J. O. Wertheim, A. L. Hernandez, M. C. Bañez Ocfemia, N. Saduvala, I. H. Hall, Using Molecular HIV Surveillance Data to Understand Transmission between Subpopulations in the United States, Journal of Acquired Immune Deficiency Syndromes (1999)doi:10.1097/QAI.0000000000000809.

[27] E. O. Romero-Severson, I. Bulla, T. Leitner, Phylogenetically resolving epidemiologic linkage, Proceedings of the National Academy of Sciences (2016) 201522930 doi:10.1073/pnas.1522930113.

[28] P. Prah, A. J. Copas, C. H. Mercer, A. Nardone, A. M. Johnson, Patterns of sexual mixing with respect to social, health and sexual characteristics among heterosexual couples in England: Analyses of probability sample survey data, Epidemiology and Infection 143 (7) (2015) 1500–1510. doi:10.1017/S0950268814002155.

[29] T. de Oliveira, A. B. M. Kharsany, T. Gräf, C. Cawood, D. Khanyile, A. Grobler, A. Puren, S. Madurai, C. Baxter, Q. A. Karim, S. S. A. Karim, Transmission networks and risk of HIV infection in KwaZulu-Natal, South Africa: A community-wide phylogenetic study, The Lancet HIV 4 (1) (2017) e41–e50. doi:10.1016/S2352-3018(16)30186-2.

[30] C. Wymant, M. Hall, O. Ratmann, D. Bonsall, T. Golubchik, M. de Cesare, A. Gall, M. Cornelissen, C. Fraser, PHYLOSCANNER: Inferring Transmission from Within-and Between-Host Pathogen Genetic Diversity, Molecular Biology and Evolution 35 (3) (2018) 719–733. doi:10.1093/molbev/msx304.

